# Serum amyloid A-dependent inflammasome activation and acute injury in a mouse model of experimental stroke

**DOI:** 10.1101/2023.06.22.546125

**Authors:** Jin Yu, Hong Zhu, Saeid Taheri, June-Yong Lee, David M. Diamond, Cheryl Kirstein, Mark S. Kindy

## Abstract

**BACKGROUND:** Serum amyloid A (SAA) proteins increase dramatically in the blood following inflammation. Recently, SAAs are increased in humans following stroke and in ischemic animal models. However, the impact of SAAs on whether this signal is critical in the ischemic brain remains unknown. Therefore, we investigated the role of SAA and SAA signaling in the ischemic brain.

**METHODS:** Wildtype and SAA deficient mice were exposed to middle cerebral artery occlusion and reperfusion, examined for the impact of infarct volumes, behavioral changes, inflammatory markers, TUNEL staining, and BBB changes. The underlying mechanisms were investigated using SAA deficient mice, transgenic mice and viral vectors.

**RESULTS:** SAA levels were significantly increase following MCAo and mice deficient in SAAs showed reduced infarct volumes and improved behavioral outcomes. SAA deficient mice showed a reduction in TUNEL staining, inflammation and decreased glial activation. Mice lacking acute phase SAAs demonstrated a reduction in expression of the NLRP3 inflammasome and SAA/NLRP3 KO mice showed improvement. Restoration of SAA expression via SAA tg mice or adenoviral expression reestablished the detrimental effects of SAA. A reduction in BBB permeability was seen in the SAA KO mice and anti-SAA antibody treatment reduced the effects on ischemic injury.

**CONCLUSIONS:** SAA signaling plays a critical role in regulating NLRP3-induced inflammation and glial activation in the ischemic brain. Blocking this signal will be a promising approach for treating ischemic stroke.

**GRAPHIC ABSTRACT:** A graphic abstract is available for this article.

---

Stroke is the second leading cause of death worldwide and is the primary cause of disability in the USA^1, 2^. The pathophysiology of cerebral ischemic injury is complex, and numerous studies have demonstrated that oxidative stress and inflammation are key in the underlying mechanisms^3, 4^. Over the years, studies have implicated various signaling pathways that are involved in the production of reactive oxygen species (ROS), mitochondrial dysfunction, and activation of apoptotic pathways^5, 6^. These include the role of NADPH oxidase and activation of the mitogen-activated protein kinase (MAPK) pathways in the cell death mechanisms and glial activation^7, 8^. In addition, the generation of inflammatory pathways and mediators as well as the role of the inflammasome in ischemic injury has been elucidated^9, 10^.

Acute-phase serum amyloid A proteins (A-SAAs) are secreted during the acute phase of inflammation^11^. SAA proteins, the most dramatic acute phase reactants, are associated with high-density lipoproteins^12^. SAA proteins are normally maintained at 1–5 μg/ml in the plasma, but during an acute phase response, levels can increase to 500–1000 μg/ml^13, 14^. SAA biosynthesis takes place primarily in the liver. SAA expression has been detected in the normal brain, but to a limited extent^15^. These proteins have several roles, including the transport of cholesterol to the liver for secretion into the bile, the recruitment of immune cells to inflammatory sites, and the induction of enzymes that degrade extracellular matrix^15–19^. A-SAAs are implicated in several chronic inflammatory diseases, such as amyloidosis, atherosclerosis, and rheumatoid arthritis^20–23^. Three acute-phase SAA isoforms have been reported in mice, called SAA1, SAA2, and SAA3^24^. During inflammation, Saa1 and Saa2 are expressed and induced principally in the liver, whereas SAA3 is induced in many distinct tissues^25^. The mouse Saa1 and Saa2 genes are regulated in liver cells by the proinflammatory cytokines IL-1β, IL-6, and TNF-α^26, 27^.

Inflammation is a response by the body to infection and injury. In the nervous system, the microglial cells react to these signals, propagate and switch to different cellular states depending on the types of signals^28–30^. NOD-like receptor proteins (NLRPs) are one of the forms of sensors in the cell membrane that are activated upon ligand binding^31–33^. The NLRP3 inflammasome complex has been associated with inflammatory responses in the brain and inhibition or genetic deletion attenuates neuroinflammation and limits the extent of injury^34–36^. Triggers that elicit NLRP3 activation result in the stimulation and release of IL-1β from the microglia cells in a pro-inflammatory retort to the injury^37, 38^. IL-1β in return can further activate inflammation in the brain as well as contribute to the degeneration and death of neuronal cells^39, 40^. Further validating that the inflammasome plays a critical role in the pathogenesis of many neurological disorders.

Previous studies have shown that SAA levels increase in the plasma following cerebral ischemia and other brain injuries, suggesting they may play a role in the outcomes^41, 42^. Inflammasome activation may also contribute to the response to injury and may be a target for intervention. To understand the mechanisms associated with SAA expression in the context of cerebral ischemia, mice deficient in the various SAAs were examined. We showed that mice deficient in the acute-phase SAA proteins had reduced inflammation and significantly less infarct volumes compared to wildtype mice. In addition, when SAAs were reintroduced by transgenics or viruses, we were able to restore the deleterious effects of the SAA proteins. These data suggest that under certain circumstances, SAAs participate in exacerbation of brain injury via enhanced inflammation mediated in part by the inflammasome complex.

## METHODS

The data that support the findings of this study are available from the corresponding author upon reasonable request.

### Study Design

All experiments were approved by the Institutional Animal Care and Use Committee of the University of South Florida (4613 and 8883) and conducted in accordance with the University of South Florida Guidelines, which are based on the National Institutes of Health’s Guide for the Care and Use of Laboratory Animals and Animal Research: Reporting of In Vivo Experiments guidelines. All procedures and histological and gene expression analyses were performed by examiners who were blinded to the experimental conditions. The number of animals was 80% powered to detect 25% changes with a 2-sided α value of 0.05. Mice that showed the following were excluded: spasm after craniotomy (n=3), spontaneous complete recovery of the middle cerebral artery during surgery (n=2), failure of the training test in the hanging wire test (n=3), and no abnormality in the hanging wire test on day 1 (n=5). None of the mice died after surgery

#### Surgical Procedures

The middle cerebral artery occlusion model was used in the study. See the Supplemental Material for details.

#### Histological Analysis

Immunohistochemistry and TTC staining were performed using 4% paraformaldehyde-fixed brain samples (Supplemental Material).

#### Real-Time Revers Transcription Polymerase Chain Reaction

mRNA was isolated from the intact or ischemic cortex collected after ischemic insult using the RNeasy Lipid Tissue Mini Kit (Qiagen, Germantown, MD) according to the manufacturer’s recommendations. See the Supplemental Material for details on further analyses.

#### Western Blotting

Western blotting and immunostaining were performed, as described previously.16 See the Supplemental Material for details.

#### Statistical Analysis

Data are expressed as the mean±SD. Data were analyzed using GraphPad Prism for Windows versions 7 & 8 (GraphPad Software, San Diego, CA). Comparisons between multiple groups were performed using 1-way ANOVA followed by Dunnett multiple comparisons test, and comparisons between 2 groups were performed using the *t* test. Differences were considered statistically significant at P<0.05.

## RESULTS

### Cerebral Ischemia/Reperfusion Injury Increases SAA Plasma Levels

We and others have previously shown that plasma SAA levels are increased following cerebral ischemia and reperfusion injury in the mouse^41, 42^. We assessed the expression of SAA in the mouse at different times of reperfusion following 1 hr of ischemia (Figure 1). Remarkably, SAA protein was expressed at high levels in the blood in ischemic/reperfused animals. As seen in the figure, SAA levels were low in control (0 hr) mice but increased dramatically at 24 hrs of reperfusion and was maintained for at least out to 120 hrs (Figure 1A). Transferrin was used as a control protein for the blood of the mice. Furthermore, in mice deficient in both *Saa1* and *Saa2* genes, showed no SAA protein at all when compared to circulating transferrin levels. Quantitation of the SAA in the blood showed a 1000-fold increase in the protein at the 24 hr time point (Figure 1B). Examination of the early time points indicated that SAA is present at least as early as 4-6 hours following ischemia and reperfusion injury (Figure 1C). Additionally, we determined that the majority of the SAA in the blood was generated by expression in the liver (Figure 1D). The different SAA isoforms are all expressed in the plasma after 1 hr of ischemia and 24 hrs of reperfusion (Figures 1E and 1F, and IEF, not shown). The marked difference in SAA expression in the liver suggests that overall release of cytokines from the brain can trigger a systemic response in plasma inflammatory proteins that could have an impact on stroke outcomes.

**Figure 1.**
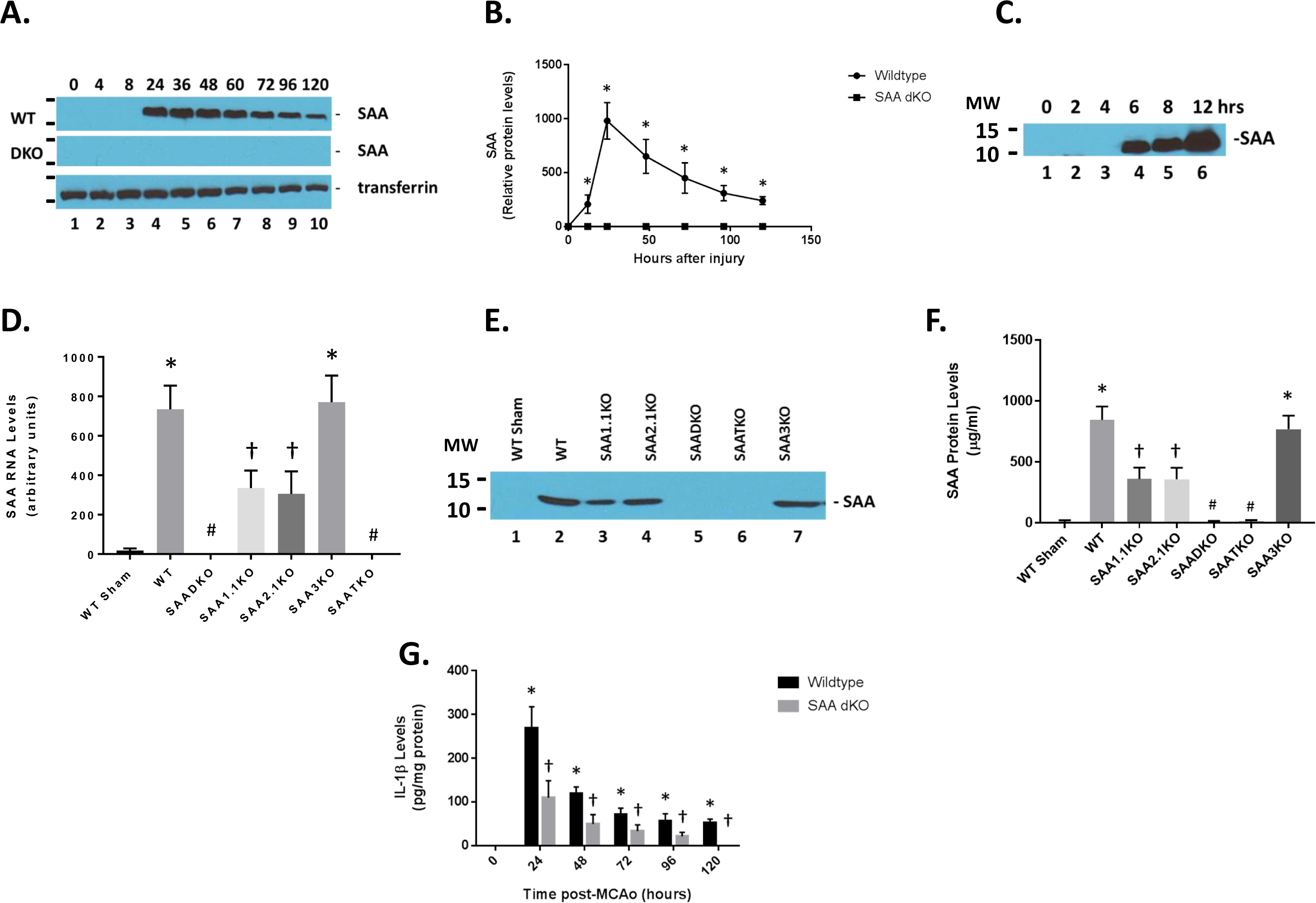
Expression of SAA proteins following ischemia and reperfusion injury. **A.** Mice were subjected to 1 hr of Ischemia and different times of reperfusion and blood was collected and subjected to western blot analysis (1/100 dilution of plasma in loading buffer). Wildtype (WT) and SAA double KO (DKO) mice were used and analyzed for SAA levels. Transferrin was used as a loading control for the plasma proteins. **B.** Graphical representation of data in A. **C.** Early time course of SAA protein expression in the blood of WT mice subjected to ischemia and reperfusion injury. **D.** Liver SAA RNA levels in the different SAA deficient mice (As determined by RT-PCR for specific mRNAs). **E.** Western blot for different SAA levels in ischemic mice. **F.** Quantitation of E. **G.** Brain IL-1β levels in the different KO were determined by ELISA. MW – molecular weight; upper band is 15 kDa, lower band is 10 kDa. Data are presented as mean ± S.E. The measurements are from 8-10 mice per group (time point). *, *p* < 0.001 compared to control (0 time point). #, *p* < 0.001 compared to WT ischemic group. †, *p* < 0.001 compared to WT sham and ischemic groups. NS=not significant. Data are considered significantly different at *p* < 0.05.

### SAADKO Mice Have Reduced IL-1β Levels

Compared to the wildtype (WT) mice, the SAADKO showed a complete obliteration of SAA expression in the plasma (Figure 1A and Figure 1E&F). However, the individual KO mice (SAA1.1 and SAA2.1) had about 50% of the total SAA expression in the WT mice. The SAA3KO mice showed similar SAA plasma levels compared to the WT animals. Finally, the SAA triple KO (SAATKO) had no detectable plasma SAA quantities.

To determine the impact of SAA expression following ischemia and reperfusion injury, mice were subjected to 1 hr of ischemia and various times of reperfusion and the brains were isolated and examined for IL-1β levels by ELISA (Figure 1G). IL-1β concentrations increased dramatically in the brain at the 24-hour time point and remained elevated for up to 120 hours after the start of reperfusion. IL-1β quantities in the SAA DKO mice were significantly reduced and decreased to undetectable levels by 120 hours. The increase in IL-1β coincides with the expression of SAA in mouse model and the data suggest that the augmentation of IL-1β is linked to SAA expression. Additionally, TUNEL analysis of the number of dead cells following ischemia and reperfusion injury in the different KO mice (and different brain regions) demonstrates the importance of the SAA proteins in neuronal cell death (Figure 2C). Finally, to validate the presence of SAAs in the brain following cerebral ischemia and reperfusion injury, following 1 hr of ischemia and 24 hrs of reperfusion, the mice were perfused to clear the blood from the cerebrovasculature and the brain was isolated for Western blot analysis. Figure 2D shows that both wildtype and individual KO mice had SAA protein present in the brain. SAADKO and SAATKO mice has little if any SAA in the brain. When SAA was expressed in the liver using adenoviral vectors, the SAA was again seen in the brain following ischemic injury. Thus, there is an impact of SAA expression following MCAO in the wildtype mice that allows for continued deterioration.

**Figure 2.**
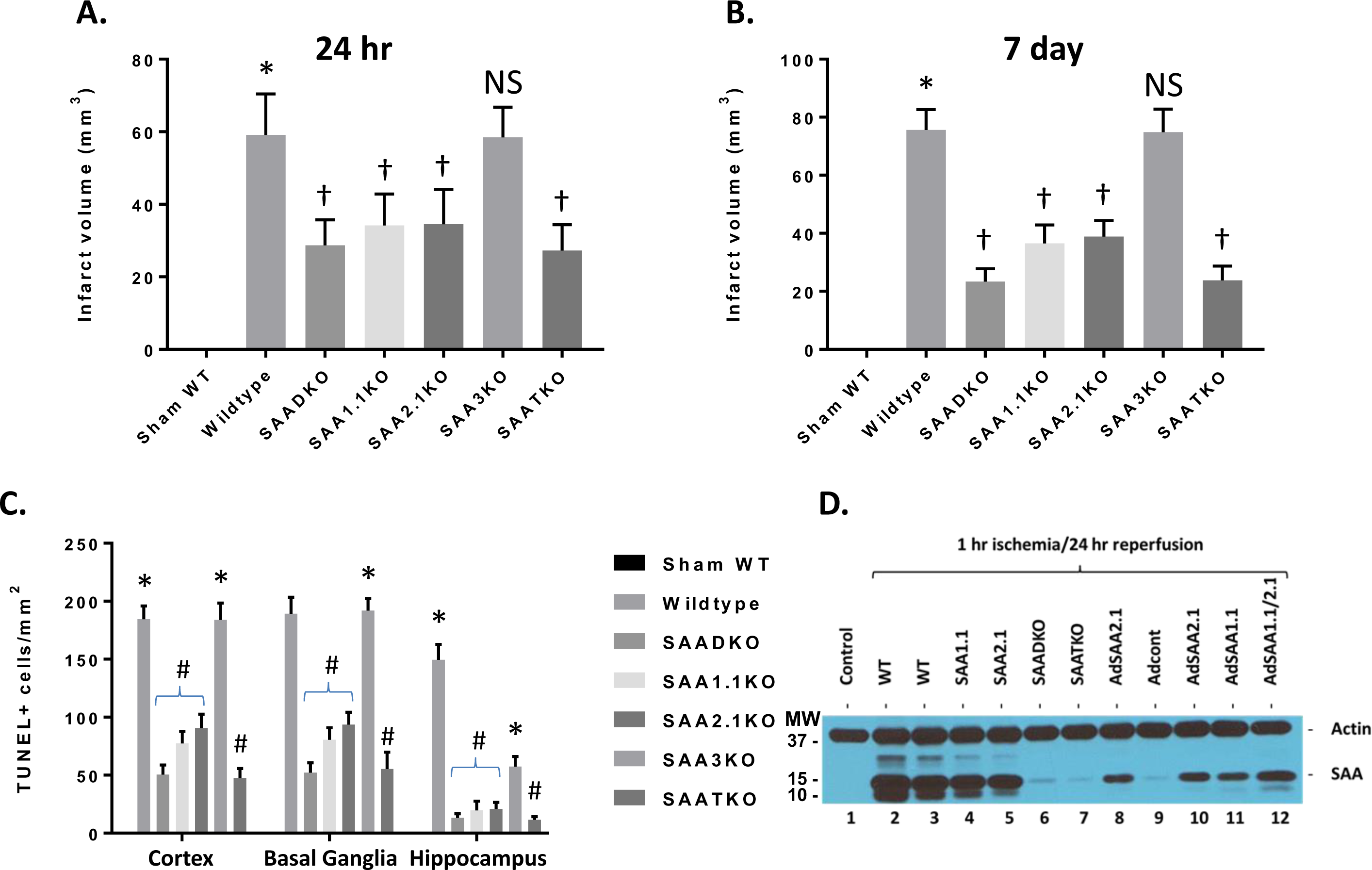
Infarct volumes and TUNEL staining in wildtype and SAA KO mice. **A.** Mice were subjected to 1 hour of ischemia and 24 hours of reperfusion and brains were analyzed for infarct volume using TTC staining. DKO – double KO, TKO – Triple KO. **B.** Mice were subjected to 1 hour of ischemia and 7 days of reperfusion and brains were analyzed for infarct volume using TTC staining. **C.** Cell death in different brain regions as analyzed by TUNEL immunostaining at 7 days after MCAO (TUNEL-positive cells per square millimeter). **D.** Western blot analysis of SAA accumulation in the brain following IRI. AdSAAs are adenoviral vectors. Data are presented as mean ± S.E. The measurements are from 8-10 mice per group. *, *p* < 0.001 compared to control. † or #, *p* < 0.01, compared to wildtype mice. NS=not significant.

### Infarct Volume is Reduced in SAA KO Mice

To further understand the role of SAA in cerebral ischemia/reperfusion injury or stroke, the different mouse strains were subjected to 1 hr of ischemia and 24 hrs of reperfusion and examined for infarct volumes (Figure 2A). Wildtype C57BL6 mice showed a typical infarct volume profile with 59.08 ± 3.27 mm^3^. While the SAADKO mice showed a significant decrease in infarct size (28.67 ± 2.03 mm^3^), the individual KO mice (SAA1.1 and SAA2.1) had a more modest reduction (34.17 ± 2.50 mm^3^ vs 34.5 ± 2.77 mm^3^, respectively). The SAA3KO mice did not show a change in the damage compared to the wildtype mice (58.42 ± 2.40 mm^3^). Finally, the SAATKO or triple knockout mice lack SAA1.1, SAA2.1 and SAA3 demonstrated an equivalent level of injury compared to the SAADKO mice (27.25 ± 2.05 mm^3^). At 7 days after the start of reperfusion, the wildtype and SAA3KO mice showed a continued increase in infarct volume (75.6 ± 2.22 mm^3^ and 74.9 ± 2.48 mm^3^, respectively), while the SAAD, SAA1.1 and SAA2.1 deficient mice showed a further reduction in or equivalent infarct size (23.4 ± 1.39 mm^3^, 36.5 ± 2.02 mm^3^ and 38.9 ± 1.74 mm^3^, respectively) (Figure 2B).

The absence of SAAs had no significant effects on blood pH or heart rate (Table 1). Compared with pre-MCAo mice, no significant changes were noted in brain temperatures or MABP for any of the treatment groups (Table 1). Comparing the pre- and post-MCAo values of blood gases revealed that MCAo showed no reduction in _P_CO_2_ or increase _P_O_2_ in KO mice.

**Table 1.** Physiological measurements in the SAA mice. Physiological measurements were taken before, during and post ischemia and reperfusion injury in the mice. Mean arterial blood pressure (MABP), heart rate (HR), body temperature (Temp) and pH were determined in the study for the different mouse strains.

|  | **Before Ischemia** |  |  |  |
| --- | --- | --- | --- | --- |
|  | MABP | HR | Temp | pH |
| Sham WT | 102.4 ± 11.5 | 419.5 ± 25.4 | 36.68 ± 0.58 | 7.33 ± 0.12 |
| I/R WT | 101.1 ± 11.2 | 418.3 ± 26.2 | 36.42 ± 0.52 | 7.34 ± 0.10 |
| I/R SAADKO | 100.5 ± 12.1 | 416.9 ± 23.5 | 36.69 ± 0.65 | 7.32 ± 0.11 |
| I/R SAA1.1KO | 101.6 ± 11.8 | 421.4 ± 24.9 | 36.73 ± 0.49 | 7.33 ± 0.13 |
| I/R SAA2.1KO | 100.9 ± 11.6 | 420.5 ± 25.2 | 36.81 ± 0.53 | 7.33 ± 0.12 |
| I/R SAA3KO | 100.4 ± 11.4 | 419.8 ± 25.5 | 36.53 ± 0.59 | 7.35 ± 0.11 |
| I/R SAATKO | 101.2 ± 12.2 | 420.3 ± 24.7 | 36.86 ± 0.55 | 7.33 ± 0.13 |
|  | **During Ischemia** |  |  |  |
|  | MABP | HR | Temp | pH |
| Sham WT | 101.2 ± 10.6 | 420.6 ± 24.7 | 36.91 ± 0.52 | 7.32 ± 0.10 |
| I/R WT | 100.5 ± 11.1 | 418.7 ± 28.1 | 36.56 ± 0.56 | 7.34 ± 0.11 |
| I/R SAADKO | 100.8 ± 11.3 | 419.4 ± 22.9 | 36.63 ± 0.61 | 7.33 ± 0.12 |
| I/R SAA1.1KO | 100.2 ± 11.5 | 417.7 ± 24.5 | 36.78 ± 0.48 | 7.35 ± 0.14 |
| I/R SAA2.1KO | 101.7 ± 11.7 | 420.5 ± 23.9 | 36.62 ± 0.54 | 7.32 ± 0.09 |
| I/R SAA3KO | 99.8 ± 10.9 | 420.3 ± 26.3 | 36.84 ± 0.58 | 7.34 ± 0.13 |
| I/R SAATKO | 100.7 ± 11.4 | 421.2 ± 25.6 | 36.89 ± 0.52 | 7.33 ± 0.12 |
|  | **Post Ischemia** |  |  |  |
|  | MABP | HR | Temp | pH |
| Sham WT | 100.5 ± 11.2 | 415.5 ± 25.4 | 36.51 ± 0.54 | 7.33 ± 0.11 |
| I/R WT | 101.3 ± 12.1 | 419.7 ± 29.3 | 36.96 ± 0.61 | 7.34 ± 0.10 |
| I/R SAADKO | 101.9 ± 10.9 | 420.4 ± 27.5 | 36.65 ± 0.55 | 7.35 ± 0.12 |
| I/R SAA1.1KO | 100.7 ± 10.7 | 421.6 ± 24.2 | 36.87 ± 0.59 | 7.32 ± 0.09 |
| I/R SAA2.1KO | 100.4 ± 11.6 | 420.1 ± 26.9 | 36.65 ± 0.62 | 7.34 ± 0.14 |
| I/R SAA3KO | 100.8 ± 11.1 | 419.7 ± 25.1 | 36.48 ± 0.47 | 7.33 ± 0.11 |
| I/R SAATKO | 100.1 ± 11.8 | 418.6 ± 23.8 | 36.73 ± 0.54 | 7.32 ± 0.13 |

We also determined cell death in the ipsilateral cortex, basal ganglia, and hippocampus at 7 days following MCAo by TUNEL staining. The SAADKO, SAATKO and individual SAA deficient mice (1.1 and 2.1) showed a significant reduction in cell death in the basal ganglia, cortex and hippocampus compared to WT and SAA3KO mice. Further, compared to individual SAA deficient mice, the SAADKO and SAATKO mice had significantly less cell death in the cortex, basal ganglia and hippocampus. There were no significant differences between SAADKO and SAATKO mice across the three brain regions (Figure 2C).

### SAA Expression on Permanent Ischemia and MCAo in the Rat

To determine the impact on SAA expression in permanent ischemia, mice were subjected to 24 hrs of ischemia with no reperfusion (Figure 3). When mice were treated with permanent ischemia, the plasma SAA levels increase in a similar fashion to ischemia and reperfusion injury (Figure 3A). The WT animals presented with the typical infarct volumes (33.4 ± 1.68 mm^3^) compared to the sham mice (Figure 3B). However, the SAA KO mice showed a reduction in infarct volume, commensurate with the level of SAA expression (SAADKO – 15.3 ± 1.45 mm^3^; SAA1.1KO – 24.5 ± 1.59 mm^3^; SAA2.1KO – 22.8 ± 1.73 mm^3^; SAA3KO – 34.3 ± 1.99 mm^3^; SAATKO – 14.6 ± 1.19 mm^3^). As seen in Figure 3A and 3D, the rat does not express a viable SAA isoform, this is due to mutations or evolutionary changes. Therefore, when rats were subjected to 2 hrs of ischemia and 24 hrs of reperfusion the infarct volume in the rat is in the absence of SAA proteins (Figure 3C, 111.4 ± 2.73 mm^3^). However, when murine SAA is delivered back using adenoviral vectors (Figure 3D), the infarct volume is significantly increased (154.8 ± 2.94 mm^3^), indicating that SAA contributes to the exacerbation of ischemic injury.

**Figure 3.**
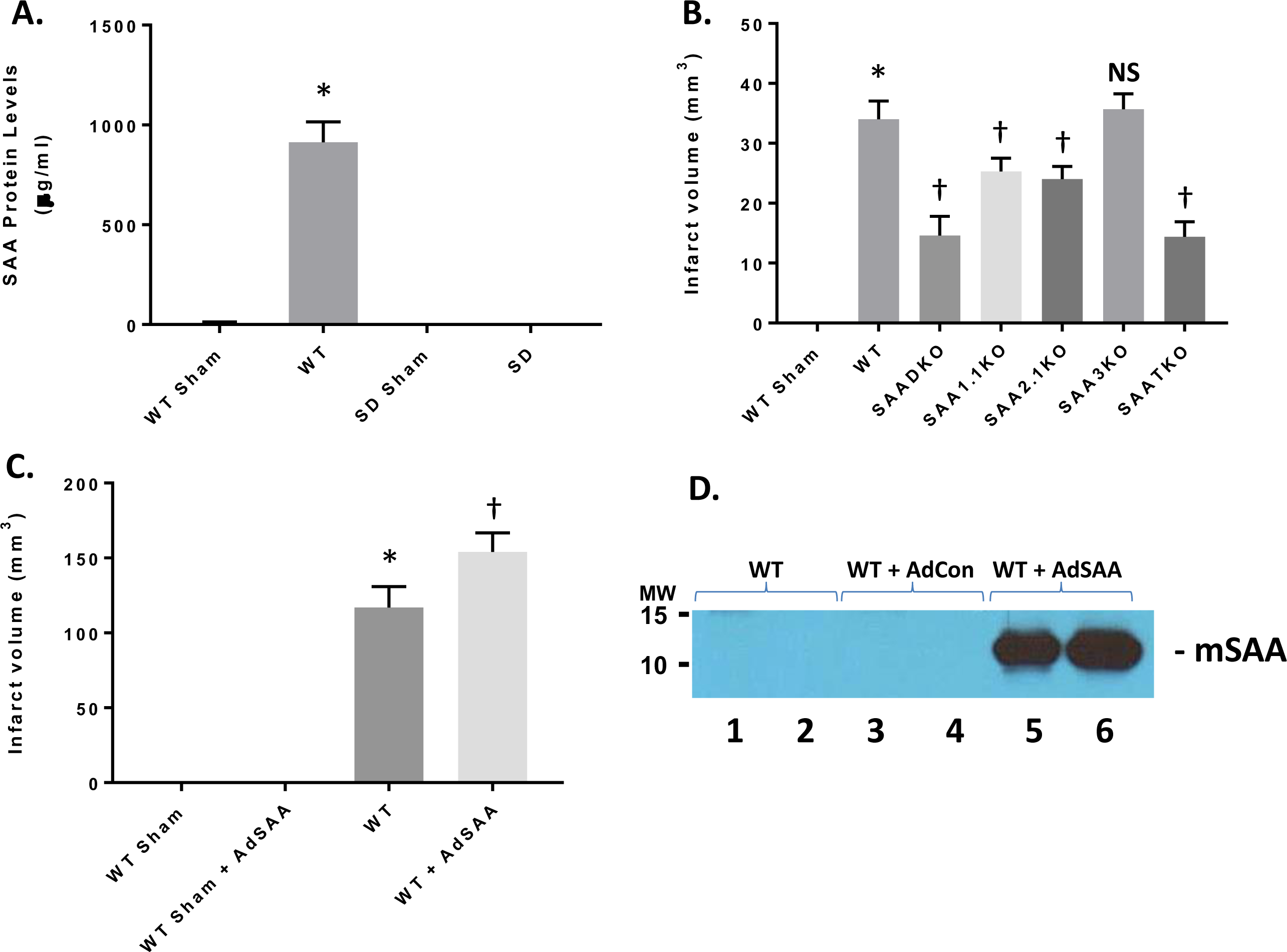
Impact of SAA in permanent ischemia in mice and I/R in in rats. **A.** Mice and rats were subjected to permanent ischemia or ischemia/reperfusion injury and examined for SAA expression. **B.** Mice (WT and SAA deficient mice) were subjected to 24 hrs of permanent ischemia and examined for infarct volumes at 24 hrs. **C.** Rats (Sprague-Dawley) were either Sham or I/R (2 hr ischemia and 24 hrs reperfusion) +/- adenoviral SAA and the infarct volume was determined at 24 hrs. **D.** Representative plasma mSAA levels from rats injected with nothing, AdCon, or AdSAA vectors, and Western blotted for mSAA expression. Data are presented as mean ± S.E. The measurements are from 8-10 mice per group. *, *p* < 0.001 compared to WT Sham. †, *p* < 0.01, compared to wildtype I/R mice. NS=not significant.

### Decreased Microglial and Astrocytic Markers in SAA KO Mice

In response to MCAo, microglia and astrocytes are activated, and microglial stimulation and astrogliosis are general hallmarks of inflammation within the brain. This process can persist for many days following a stroke and may contribute to the disease process. Microglial activation and astrogliosis was assessed in the brain at 7 days post-MCAO by immunohistochemical detection of Iba-1 and GFAP markers in the cortex, respectively (Figure 4). Compared to WT sham animals, there was a significant reduction in both activated microglia (Figure 4A & C) and reactive astrocytes (Figure 4B & D) in the SAADKO, SAATKO and to a lesser extent in the individual SAA deficient mice. In contrast, there was no significant difference in the detection of either marker in SAA3KO mice compared to WT. No differences were observed in the contralateral hemispheres between the different groups. These data indicate that SAA deficiency reduce glial cell activation after MCAO.

**Figure 4.**
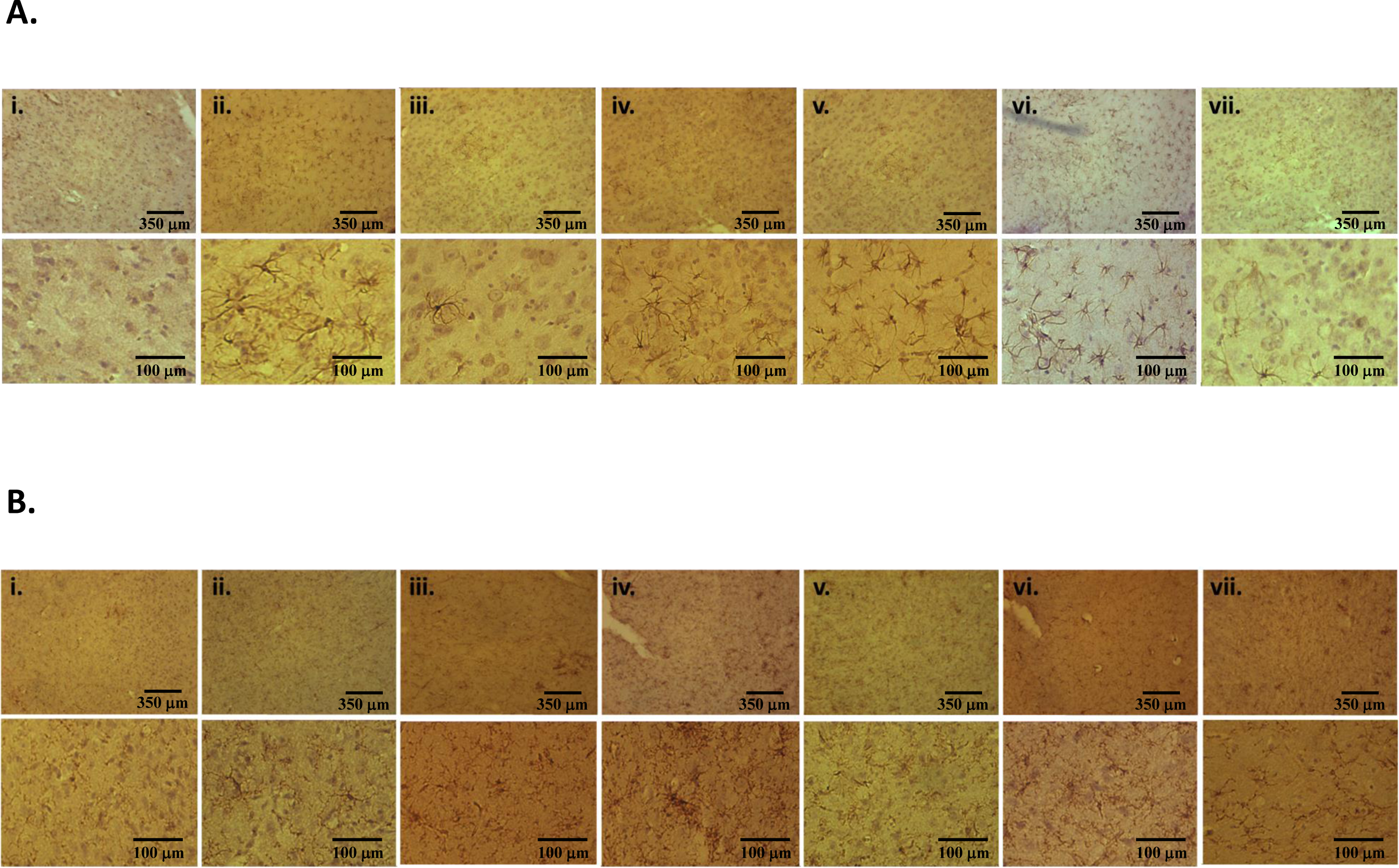
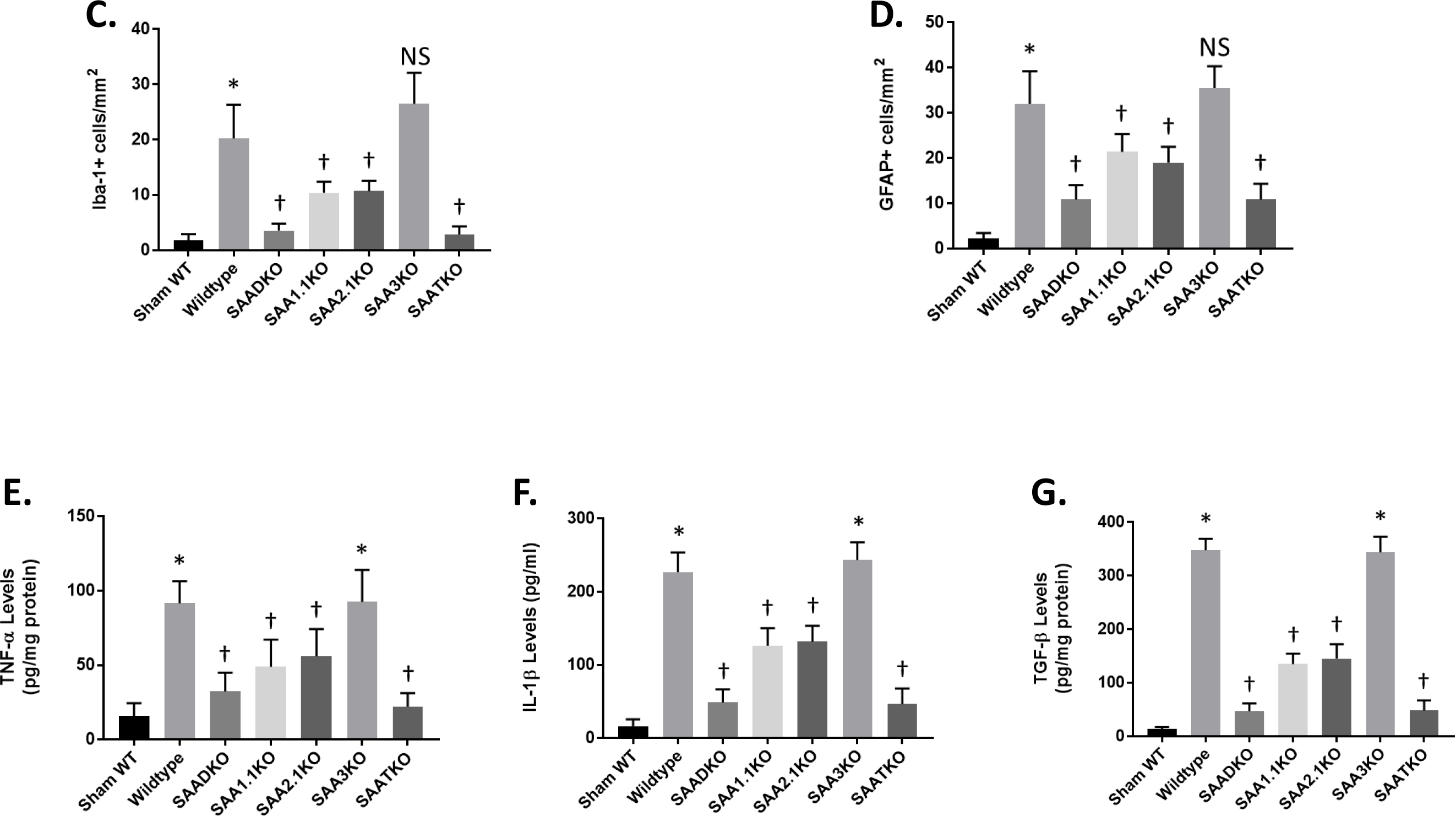
Microglial, astrocytic and inflammatory activation following IRI in SAA deficient mice. **A.** Brains of mice from 1 hr ischemia and 24 hr reperfusion were sectioned and stained with iba-1 antibody. i - Sham WT; ii - WT; iii – SAADKO; iv – SAA1.1KO; v – SAA2.1KO; vi – SAA3KO; vii – SAATKO (top panel 10X, bottom panel 40X). **B.** Brains of mice from 1 hr ischemia and 24 hr reperfusion stained with gfap antibody. Order is the same as A (top panel 10X, bottom panel 40X). **C.** Quantification of data in A. **D.** Quantification of data in **B**. Mice were subjected to 1 hr ischemia and 24 hrs of reperfusion and the brain isolated for cytokine analysis. **E.** Brains were analyzed for TNF-α levels. **F.** Brains were analyzed for IL-1β levels. **G.** Brains were analyzed for TGF-β levels. Data are presented as mean ± S.E. The measurements are from 8-10 mice per group. *, *p* < 0.001 compared to control. †, *p* < 0.001, compared to wildtype mice or other groups. NS=not significant.

To determine the impact of the SAA deficiency on neuroinflammation in the MCAo mouse brain, we determined the expression of inflammatory markers. We measured the levels of specific cytokines namely tumor necrosis factor-α (TNF-α), interleukin-1β (IL-1β), and transforming growth factor-β (TGF-β) 24 hrs after reperfusion (Figure 4E-G). The data showed that when SAAs were removed from the equation, there was a significant reduction in TNF-α (E), IL-1β (F), and TGF-β (G) levels.

### Attenuated Behavioral Changes are Associated with SAA Deficiency

To better understand the impact of SAA deficiency on stroke outcomes, we measured neurological severity score via locomotor activity in an open field activity monitor (Figure 5). There was a significant difference in neurological severity score (NSS) between the groups, WT and SAA3KO mice started out with a higher score that did improve slightly over time (Figure 5A). While the SAADKO, SAA1.1KO, SAA2KO, and SAATKO started off with a better (lower) score and improved to a greater degree than the WT mice. In the locomotor activity chambers, the SAA deficient mice showed significantly increased locomotor activity (total distance and # of movements) compared to the WT and SAA3KO mice (Figure 5B & C). Studies have suggested that anxiety may impact the outcomes of the behavioral testing in the mice, therefore, we evaluated the anxiety levels across the groups by measuring the percentage of time spent at the center of the open field. We found that MCAo did increase the anxiety in the WT and SAA3KO mice significantly compared to the other SAA deficient mice which reduced the anxiety levels in the different groups on days 3 and 7 post-MCAO (Figure 5D). The data implicate SAA as a factor that is involved in the evolution of the injury to the brain following MCAo and might be a potential target for therapeutic intervention.

**Figure 5.**
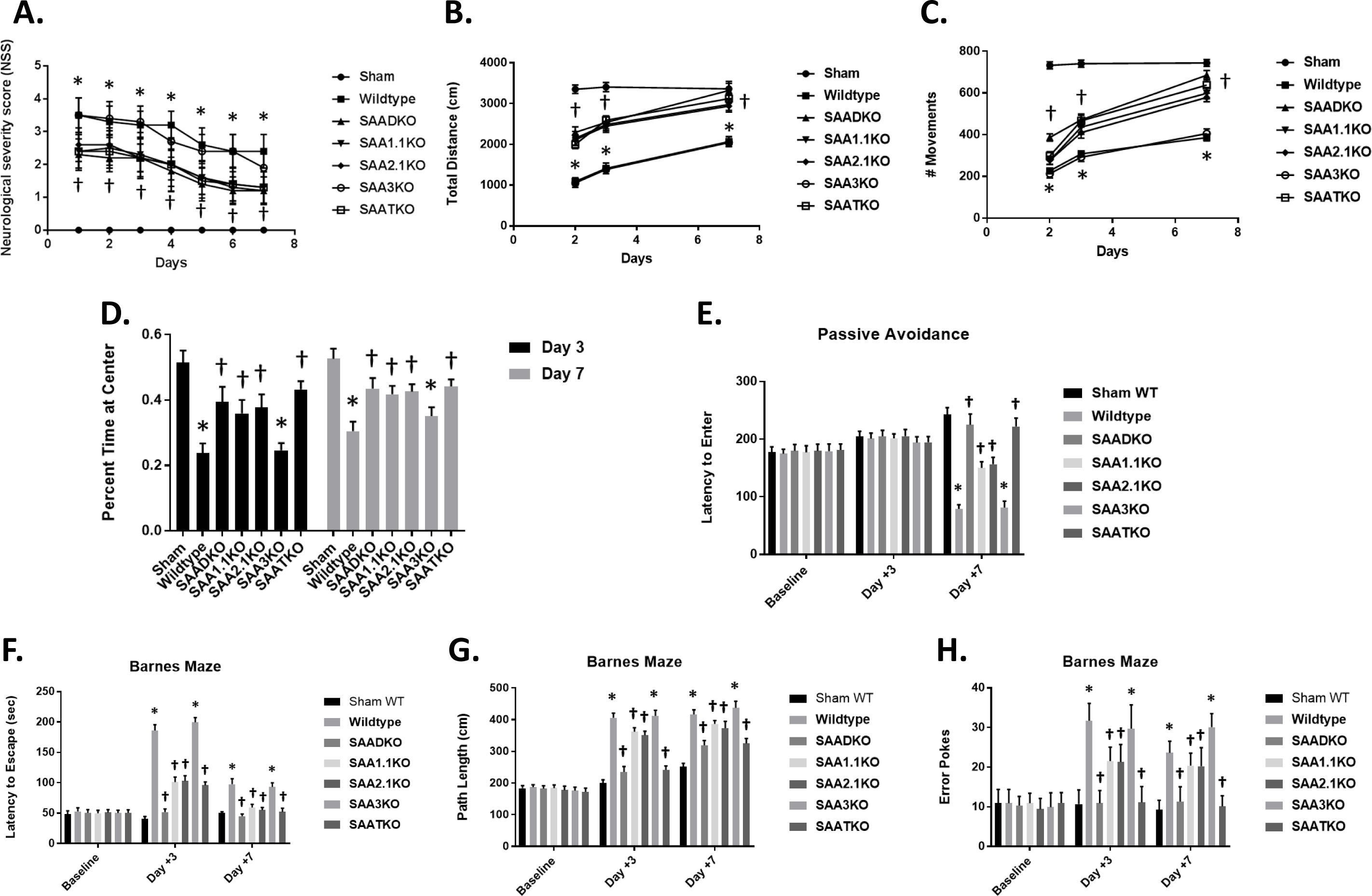
Effect of SAA deficiency on behavioral deficit and locomotor function after MCAo. **A.** Neurological deficit scores (0–4) within the first week after MCAo. **B & C.** Locomotor activity measured as total distance moved **(B)** or number of movements **(C)**, using an open field activity monitor. Determinations were made 2, 3, and 7 days after MCAO. **D.** Anxiety assessment across the different groups measured as percent time spent at the center of an open field at 3 and 7 days. **E.** SAA deficiency improves performance on passive avoidance task 7 days after MCAo. Shown is latency to enter a dark box associated with a shock. Mice were given a trial to associate a shock with the dark side of an apparatus, and latency to enter dark side was evaluated on days 3 and 7 post-MCAo. **F**–**H**, SAA deficiency reduces spatial memory deficits in Barnes maze task in the subacute phase after MCAo. Mice were trained on the maze for 5 days before surgery and then tested on days 3 and 7 after MCAo for latency to escape **(F)**, path length **(G)**, and number of error pokes **(H)**. Data are presented as mean ± S.E. The measurements are from 8-10 mice per group. *, *p* < 0.001 compared to control. †, *p* < 0.01, compared to wildtype mice.

### Lack of SAA improves behavior in subacute phase after stroke

Based on the significant sparing of tissue resulting from SAA deficiency, we performed a more comprehensive investigation of cognitive outcome by investigation of spatial reference memory and avoidance learning using Barnes maze and passive avoidance tasks. In the passive avoidance task, there were no significant differences between WT and SAA KO mice at either baseline or 3 days post-MCAo, with mice from all the groups recording latency to enter times similar to the times that were recorded at baseline (Figure 5E). However, on day 7 post-MCAo, there was a significant reduction in performance of the WT group, whereas the performance of SAA deficient mice was significantly different to the baseline and day 3.

In the Barnes Maze task 3 days post-MCAo, the performance of WT mice was significantly worse, as demonstrated by an increase in latency to escape, path length, and number of error pokes compared to baseline. On day 7 post-MCAo, the performance in the WT group was improved, but was still significantly poorer compared to baseline (p<0.001). In contrast, the SAA deficient mice displayed no or moderate impairment on the Barnes maze performance at either 3 or 7 days post-MCAo compared to baseline (Figures 5F-H). Consequently, compared to WT, the SAA deficient mice had significantly better spatial reference memory through the subacute phase after ischemic stroke.

### SAA Inflammatory Processes are Mediated by NLRP3

Previous studies have implicated SAAs action via the node-like receptor protein 3 (NLRP3) and activation of the inflammasome^41^. To determine the impact of SAA in the MCAo model, WT, SAADKO, NLRP3KO, and NLRP3/SAADKO mice were examined for the expression of NLRP3 following 1 hour of ischemia and 24 hours of reperfusion (Figure 6A). As seen in the figure, in WT mice, NLRP3 is dramatically increased compared to the non-ischemic mice. However, in the NLRP3 deficient mice, obviously NLRP3 is non-existent, while in the SAADKO, the NLRP3 is significantly reduced. When assessing the infarct volume in the mice, the NLRP3 deficient mice have reduced infarct volumes (as previously shown) and is significantly lower than the SAADKO mice (Figure 6B). In addition, when the NLRP3 and SAAD deficient mice were crossed together, the infarct volume was comparable to the NLRP3 deficient mice alone, suggesting that the majority of the SAA effect is mediated through the inflammasome.

**Figure 6.**
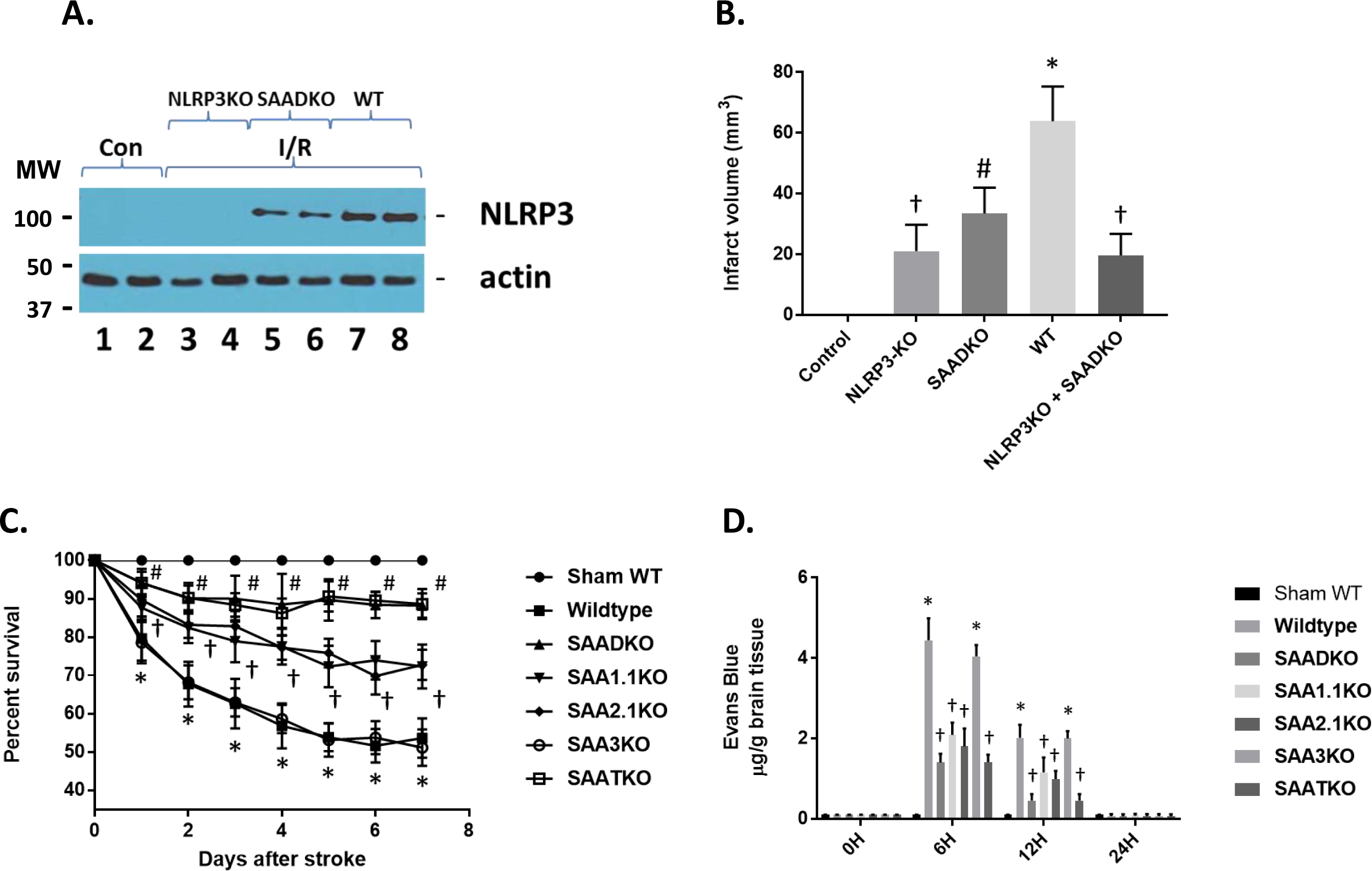
NLRP3 levels following cerebral ischemia and reperfusion injury. **A.** Wildtype (WT), SAA double knockout (SAADKO) and NLRP3 KO mice were subjected to 1 hr ischemia and 24 hrs of reperfusion and the brain isolated for protein analysis. Protein was subjected to Western blot analysis for NLRP3 and actin. Con (control); I/R (ischemia and reperfusion). **B.** Graphical representation of data in A, and additional strains. NLRP3 + SAADKO are KO for NLRP3, SAA1.1 and SAA2.1. **C.** Kaplan-Meier survival analysis of wildtype, and different SAA deficient mice over a 7-day period post-MCAO. **D.** Mice were subjected to 1 hr ischemia and different times of reperfusion to 24 hr and the brain isolated for Evans blue measurement. Mice were injected via the tail via 30 minutes prior to euthanasia. Data are presented as mean ± S.E. The measurements are from 8-10 mice per group. †, *p* < 0.001 compared to control (non-ischemic). #, *p* < 0.001 compared to control and NLRP3KO. *, *p* < 0.001, compared to control and all other groups. †, *p* < 0.001 compared to other groups.

Complications from a stroke are a major issue in patients following strokes and can increase the risk for death and disability. Since SAA levels are increased not only in response to injury but as a result of infection and suppression of the immune system, we examined the impact of SAA deficiencies on the survival rate in our model system (Figure 6C). MCAo in the WT and SAA3KO mice showed a significant effect on survival in that by 7 days, almost 50% of the animals did not survive. However, mice deficient in SAA1.1, SAA2.1 or both showed a significant improvement in survival out to 7 days, with the SAADKO and SAATKO demonstrating a greater survival rate than the individual SAAs. These data indicate that treatment to reduce or impede the role of SAA in MCAo might reduce not only the infarct volume but decrease susceptibility to secondary complications seen in experimental ischemic stroke.

To determine the impact of SAAs on BBB dysfunction post-stroke, C57/BL6 male mice underwent MCAo and were injected with 2% Evans blue as a marker for BBB penetration and permeability 30 minutes prior to euthanization (Figure 6D). Evans Blue penetrated the BBB at 6 hours and 12 hours then returned to normal levels. Quantification of Evans Blue extravasation into the hemispheres revealed a 4-fold increase at 6 hours, post-stroke in the right hemisphere of the brain where the occlusion occurred. There was no significant Evans blue extravasation observed at 24 hours post-stroke. The SAA deficient mice have a reduced BBB penetration of Evan’s blue at both the 6- and 12-hour time points. SAA3KO mice on the other hand, showed no difference in the Evans blue extravasation compared to the WT animals. These data suggest that BBB opening is partially dependent on the presence of SAA.

### Restoration of SAA in SAA KO Mice Reestablishes Activity

In order to determine the direct role of SAA in the pathogenesis of stroke, we used two different approaches. We obtained the mouse SAA transgenic (mSAAtg) mouse with the transgene under an inducible promoter with doxycycline (dox)(Figure 7A). When the mice were given dox (2 mg/ml) in their drinking water, the amount of SAA in the blood increased significantly to well over 1000 mg/L. As long as the animals are on the dox the SAA levels will stay elevated. Previous, we have developed adenoviral vectors that express the different mouse SAA (1.1 and 2.1) proteins. When the adenoviruses were injected into the mice, the expression levels were similar to an induction by LPS and the transgenic mouse (Figure 7B). To validate the roll of SAA in stroke, the mSAAtg mouse was crossed onto the SAADKO mouse, and the SAADKO were injected with SAA adenoviruses. As seen previously, the SAADKO mice had reduced infarct volumes compared to the WT animals (Figure 7C). However, when SAA was expressed via transgene or viral vector, the infarct volumes returned to WT levels. These studies demonstrate that SAA play a critical role in the outcomes from stroke.

**Figure 7.**
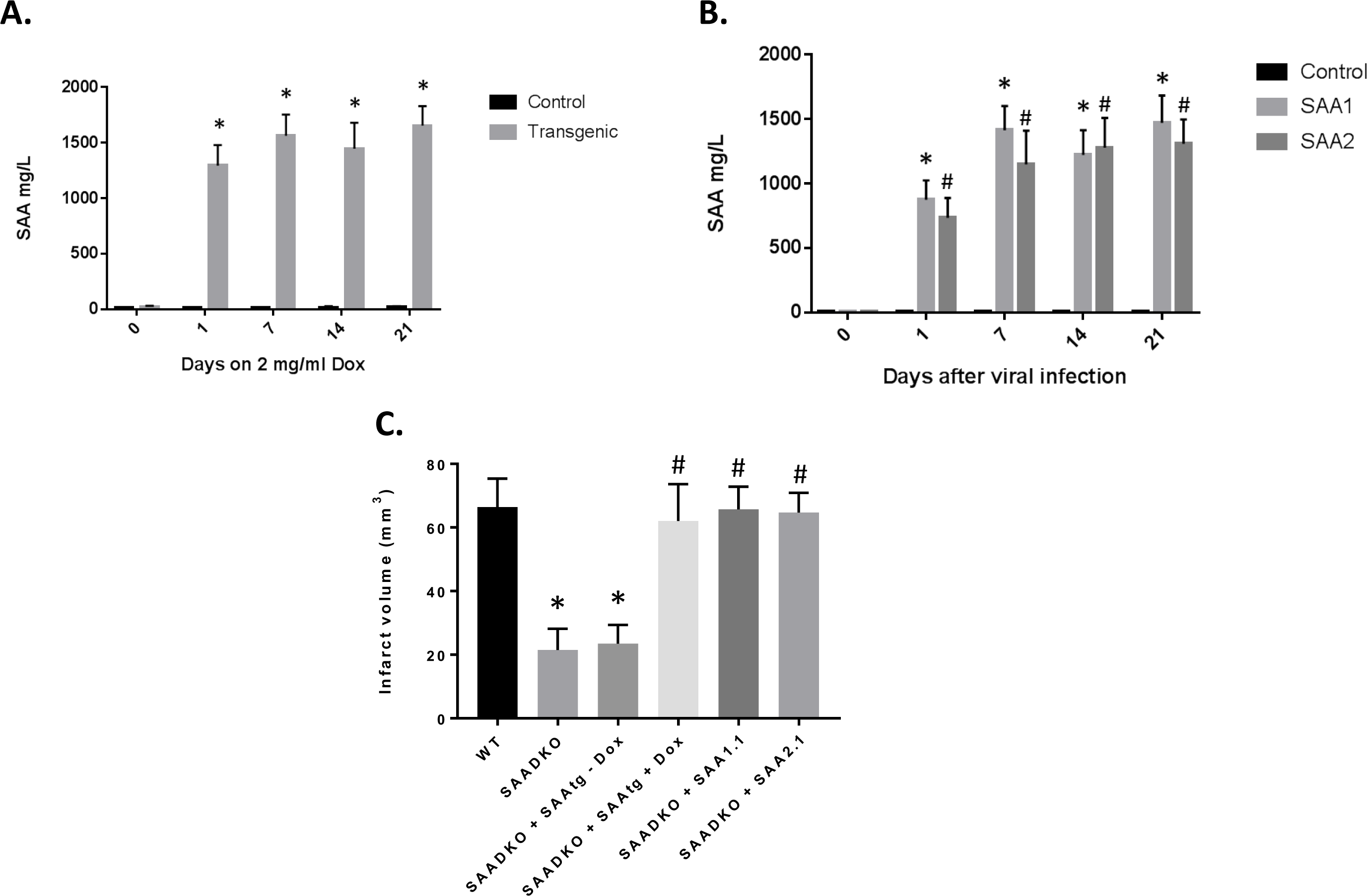
Expression of SAA via transgene or adenoviral vectors restores the detrimental effects of the proteins. **A.** Mouse SAA transgenic mice were crossed onto SAADKO background and subjected to Dox (2 mg/ml) in drinking water and plasma was examined for SAA protein levels by ELISA. **B.** SAADKO mice were injected with adenoviruses containing the SAA1.2 or SAA2.1 cDNAs and plasma was examined for SAA protein levels by ELISA. **C.** Mice were examined for infarct volumes in the presence of SAA mediated through the activated transgene or viral vectors. Data are presented as mean ± S.E. The measurements are from 8-10 mice per group. *, *p* < 0.001 compared to control group. #, *p* < 0.001 compared to control group or to treated groups.

### Protection from IRI with SAA Neutralizing Antibodies

We tested the therapeutic potential of interfering with liver SAA expression in the MCAo mouse model that develops increased SAA levels, infarct volume and inflammation following cerebral ischemia and reperfusion injury. We administered the specific mouse anti-SAA blocking antibody (25 mg/kg body weight via intraperitoneal injection) beginning at the time of reperfusion. After 24 hrs of reperfusion, NSS and IL-1β and infarct volume were determined (Figure 8). As seen in the figure, the blocking or neutralizing antibodies (nAb) showed a reduction in infarct volume in the WT mice after MCAo, while the control antibody (Ab) had not effect (Figure 8A). In addition, in the SAADKO mice, the nAb had no improvement over the lack of SAA in the MCAo model. Figure 8B shows that the nAb reduced the IL-1β levels in the brain following MCAo when compared to control and Ab treatment, while having little effect in the SAADKO mice. Finally, we showed that the SAA blocking Abs reduced the NSS and behavioral detriment seen in the MCAo model (Figure 8C). These data indicate that blocking SAA inhibits inflammation in the brain following MCAo and attenuates the development of the infarct volume and neurological deficits seen in a well-established animal model of MCAo.

**Figure 8.**
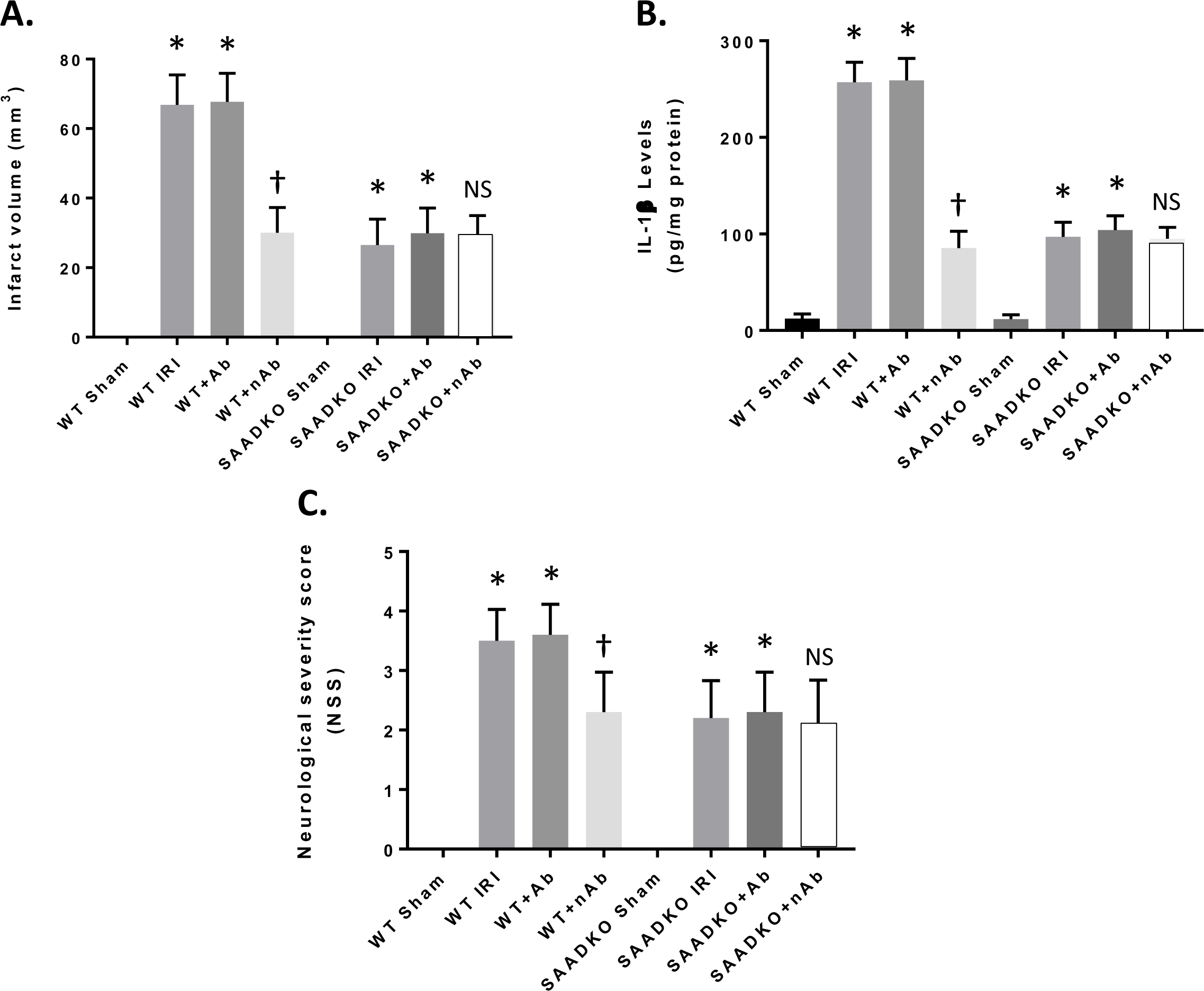
Reduced Effects of SAA with neutralizing antibodies. Mice were subjected to 1 hr ischemia and were injected with anti-mouse SAA antibodies (neutralizing) at 50 mg/kg (i.p.) and reperfusion to 24 hr. Mice were examined for infarct volume (A), IL-1β expression (B) and neurological severity score (C). Data are presented as mean ± S.E. The measurements are from 10 mice per group. *, *p* < 0.001 compared to sham group. †, *p* < 0.001 compared to other groups. NS, not significant compared to other groups.

## Discussion

Our data show that SAAs are a critical constituent of the inflammatory mechanism that contributes to the pathogenesis of stroke^41–43^. These data confirm the functional significance of SAA expression in the brain and identifies SAAs as decisive components that regulate the inflammasome pathway^44^. Thus, the SAA-inflammasome axis appears to be a major route mediating inflammation in the brain via the systemic circulation which is in contrast to microglia to neuron interactions that dominate most neurological disorders^45, 46^. It was of major interest that the SAA proteins were dramatically elevated in ischemia and reperfusion injury in the mouse model (as well as seen in stroke patients). It is likely that the SAA proteins, which are for the most part elaborated from the liver, are increased in response to the release of cytokines (IL-1β, IL-6, TNF-α, etc.) from the brain following ischemic injury^47^. Release into the blood stream and transport to the liver, will trigger the increase in SAA expression which in turn would be transported to the brain via the blood either on high density lipoprotein (HDL) particles or other particles or even as non-lipidated entities^48, 49^. Once in the brain circulation, either through breakdown of the BBB or transport across the BBB would allow for access to the neurons and glial cells^41, 45, 50^. This in turn would elicit the responses seen in the present study. Therefore, inhibition of SAA expression might provide a mechanism to down-regulate the SAA-inflammsome axis and afford protection following ischemia and reperfusion injury.

Compared to the WT mice, the SAA deficient mice showed a significant decrease in neuronal cell death following acute ischemic injury as well as in ischemia and reperfusion injury^41^. The studies presented here are carried out using constitutive knockout mice, and SAA expression are deleted from all organs, so we cannot rule out the potential impact of SAA expression in glial cells or neurons on the outcomes^51, 52^. However, several pieces of evidence strongly point to the role of liver expressed SAAs in the pathogenesis of stroke. First, ischemia and reperfusion injury results in a significant increase in SAA expression from the liver as determined by both mRNA and protein levels. Second, SAA KO mice show a significant reduction in neuronal injury in the ischemia and reperfusion animals, and even though SAAs have been shown to be expressed in the brain, the expression levels are quite low. Third, restoration of liver specific expression of SAAs via transgenic mice and adenoviral vectors implicates the majority of the neuronal injury to this specific level of articulation. Fourth, inhibition of SAAs via specific immunotherapy can provide protection to the brain following ischemia and reperfusion injury, although it could be argued that this could interfere with brain expression as well. Finally, in the rat model of ischemia and reperfusion injury, great times of ischemia are required to generate similar levels of injury as in the mouse, and this may be attributed to the lack of SAA expression in the rat. We validated these results by expressing mSAAs in the rat and were able to restore the efficacy of the SAA proteins in exacerbated neuronal injury, suggesting that SAA plays a critical role in brain injury. These studies indicate that inhibition of SAA expression following a stroke, via immunotherapy, antisense oligonucleotides or specific inhibitors might afford protection to the brain^53, 54^.

Our data suggests that targeting SAAs might be a potential therapeutic approach to reduce the impact of stroke and to extend the window of opportunity for stroke recovery. Currently, there are not therapeutic treatments for ischemic stroke, although numerous neuroprotectants, anti-inflammatory agents, hypothermia, etc., have shown to be effective in various animal models and even in phase I and II studies^55, 56^. Patients with occlusion of the major arteries of the brain are often treated with intravenous tissue-type plasminogen activator (IV tPA) thrombosis and/or mechanical thrombectomy (MTE)^57^. However, the time period for the delivery of these agents or mechanical disruption is limited and time is of the essence^58^. The damage to the brain continues from the core into the penumbra region and beyond that results in poor clinical outcomes. By inhibiting or reducing SAA expression immediately following, this stroke may attenuate the damage and allow for time for recanalization and recovery from the progression in the neurological deficits.

Moreover, our data show that neuroprotection will be efficacious in the stage of secondary brain injury within the context of ischemia and reperfusion injury and that be blocking SAA entry into the brain might provide a powerful approach as an adjuvant therapy in acute stroke to reduce the damage and improve outcomes^41^. Complications that occur following ischemia and reperfusion injury are hemorrhagic transformation, bleeding and infections that that worsen outcomes and lead to disability and death^59^. A number of therapeutic approaches have met their demise due to increased complications associated with bleeding. This includes tissue plasminogen activator (tPA) even when delivered under the prescribed conditions. In contrast, the SAA deficient mice had reduced bleeding in the brain following ischemia and reperfusion injury, as well as bleeding times were slightly reduced in the mice. Finally, the SAA deficient mice showed improved long-term survival compared to WT mice suggesting a reduction in infection related outcomes. Therefore, inhibition of SAA might provide an opportunistic therapeutic approach for acute stroke patients by reducing inflammation, lesion volumes, bleeding and response to infection.

In addition to stroke, inhibition of SAAs might be a novel treatment for other neurological and neurodegenerative disorders in which disturbances in SAA homeostasis comprise a major pathogenic feature^60, 61^. The recognition of SAAs as a significant constituent of stroke and responds to cytokine triggers released by the brain, provides a more viable target for pharmacological intervention. Given that the SAA deficient mice are healthy in their appearance and are unaffected in their general cognitive behavior or other physiological aspects, the inhibition of SAAs might be a rather safe approach to treat these disorders. In addition, the lack of side effects might prove beneficial in the treatment paradigm.

In conclusion, our findings establish SAAs as a major mediator of inflammation in the brain that plays an important role in ischemia/reperfusion injury. These findings may provide evidence for the development of therapeutics approaches using anti-SAA treatments in stroke.

## Nonstandard Abbreviations and Acronyms

BBB: blood brain barrier
IL1β: interleukin-1β
IL6: interleukin-6
SAA: serum amyloid A
TNF-α: tumor necrosis factor-α

## Acknowledgements

We thank Dr. Frederick DeBeer and for the SAA1.1, 2.1 and DKO mice, Dr. Alan Chait for the SAA3 KO mice, Dr. Dan Littman for the SAATKO mice, Dr. Doug Golenbock for the NLRP3 KO mice, and Dr. Paul Simons for the SAA tg mice.

## Author Contributions

Designed the experiments: J.Y. and M.S.K. Conducted the experiments: J.Y., H.Z., S.T., M.S.K. Analyzed and interpreted the data: J.Y., S.T., D.M.D., C.K., J.Y.L., and M.S.K. Wrote the manuscript: M.S.K. All authors edited and approved the manuscript.

## Supplemental Material

Supplemental Methods References 62-77

## Methods

### Animals

Adult C57BL/6 male and female mice (30-35 gms, 5-6 months of age), SAA1.1KO, SAA2.1KO, SAA3KO, SAA-double KO (SAADKO), SAA-triple KO (SAATKO) (all on a C57BL/6 background) were bred as needed as the University of South Florida^62–64^. Five to 6 month old animals were used to better represent individuals that have strokes. Normally 10-12 week old animals are used which are too young. Animals were randomly assigned to different groups, and the individuals who performed the animal studies were blinded to the different strains and groups. The behavioral testing experimenters were also blinded to the different groups. All the studies followed the STAIR and ARRIVE guidelines for preclinical studies^65, 66^. The experimental procedures were approved by the Institutional Animal Care and Use Committee of the University of South Florida.

### Ischemic Injury and Infarct Volumes

Transient (60 minutes) focal ischemia was induced by suture occlusion of the middle cerebral artery (MCAo), as described previously^67, 68^. Mice were anesthetized using a mixture of 1.5% isoflurane, 70% N_2_O, and 28.5% O_2_. After the ischemia (1 hr), reperfusion was started and continued until the end of the study. Body temperatures were maintained at 37◦C by a water-jacketed heating pad and anal thermometer. Transcranical laser doppler was used to monitor any changes in the cerebral blood flow. Mice were included in the study when the blood flow dropped to below 20% during ischemia and returned to 90% during reperfusion, of the normal value for data analysis. After the indicated times of ischemia and reperfusion injury (IRI), mice were euthanized, and the brains were removed for analysis. Coronal sections at 1-2 mm intervals were prepared and stained with 2% 2,3,5-triphenyltetrazolium chloride (vital dye). Infarct volumes were calculated by summing the infarcted areas (pale) of all sections and multiplying by the thickness of the sections.

### Measurement of cerebral blood flow

Regional cerebral blood flow (rCBF) was analyzed by laser Doppler flowmetry every 30 minutes over the period 1 hour before to 6 hours after MCAo^69^. Mice were anesthetized with isoflurane (1.5% in 70%/28.5% NO_2_/O_2_) and a 2-mm hole was drilled in the skull, with the probe was positioned at 0.1 mm above the dura over the cortical surface. In the hemisphere ipsilateral to the occlusion, coordinates were as follows: point A, 1 mm posterior to the bregma and 5.4 mm lateral to the midline; point B, 1 mm posterior to the bregma and 2.1 mm lateral to the midline; point C, 1 mm anterior to the bregma and 3.4 mm lateral to the midline. The mean values of rCBF were measured before MCAO as baseline, and the data thereafter were expressed as percentages of the baseline value. The rCBF data in the present report were taken from reference point A.

### TUNEL Staining

Following ischemia and reperfusion injury, the terminal deoxynucleotidyl transferase (TdT)-mediated dUTP-digoxygenin nick-end labeling (TUNEL) technique was used to determine the number of apoptotic cells (ApopTag® Peroxidase *In Situ* Apoptosis Detection Kit, Chemicon International, Inc., Temecula, CA, USA)^70^. The preparation of the cryosectioned tissue was the same as described above. TUNEL assays were executed according to the manufacturer’s instruction. In brief, the sections were fixed in 1% paraformaldehyde for 10 min at room temperature, and then post-fixed in pre-cooled ethanol: acetic acid (2:1) for 5 min at − 20 °C. The endogenous peroxidase enzymes were quenched in 3.0% H_2_O_2_ for 5 min at room temperature (RT). Subsequently, the sections were incubated with buffer for at least 30 s at RT, which was followed by incubation with TdT in a humidified chamber at 37°C for at least 1 h. The reaction was terminated by adding the stop/wash buffer for 10 min at RT. Sections were then incubated with anti-digoxigenin peroxidase conjugate in a humidified chamber for 30 min at RT and, developed with 3, 3′-diaminobenzidine (DAB) for 5 min at RT. Sections were then counterstained with 0.5% methyl green and cover-slipped. Sections from 8 to 10 separate animals were used to assess the number of TUNEL-positive cells.

### Neurological outcomes

Neurological deficits were determined following MCAo by blinded observers using a five-tiered scoring system, as previously described^71, 72^. Locomotor activity was automatically quantified using the open field activity monitor (Any Maze, San Diego Instruments). Mice were placed in a random corner and allowed to acclimate for 10 min prior to a 60-min testing period. External noise, lights, and other stimuli were minimized to reduce bias. A number of measures were automatically obtained during the task including total distance, number of movements, time spent at periphery, and time spent at the center of the enclosure. Activity readings acquired prior to sham procedure were used to establish baseline activities. The duration that the animal spent at the periphery vs. the center was used to assess anxiety level during the task.

### Behavioral Testing

Animals were tested for their performance on both Barnes maze and passive avoidance tasks^72^. To evaluate spatial reference memory, mice were educated on Barnes maze for 5 days before surgery, as previously described, then tested again on days 3 and 7 after the start of reperfusion for the time needed to escape into the hole, the number of error pokes, and the length of the animal’s path prior to escape. An automated passive avoidance apparatus (San Diego Instruments) was used to assess avoidance learning with automated sensing and shock systems (Gemini, San Diego Instruments). The instrument included a double compartment chamber with one lit and one dark compartment. Mice were allowed to explore the chamber for 5 min for orientation. Following habituation, the mice were given one trial where a shock is associated with the dark side, allowed 48 h of rest, and then tested for retention measured as latency to enter the dark side. Testing was repeated on days 3 and 7 post-reperfusion with no shock delivered during test phase.

### Immunohistochemical Analysis

Immunohistochemical staining was conducted on 8-μm paraffin sections and assessed by a blinded observer by light microscopy (Olympus BX61)^72^. Following antigen retrieval (IHC World), the following primary antibodies were used: anti-ionized calcium-binding adaptor molecule 1 (Iba-1, 1:250; Abcam), and anti-mouse glial fibrillary acidic protein (GFAP, 1:1000; Dako). Primary antibodies were detected with ImmPress-HRP kit and NovaRed peroxidase chromagen (Vector Laboratories), and primary antibodies were omitted for negative controls. Terminal deoxynucleotidyl transferase dUTP nick end labeling (TUNEL) staining was performed using ApopTag Peroxidase Staining Kit (Millipore) per manufacturer’s instructions.

### RNA Isolation and Analysis

Total RNA was isolated from mouse livers using TRIzol Reagent (Invitrogen/ThermoFisher) according to the manufacturer’s instructions. RNA samples (10 µg) were treated with DNase 1 (TURBO DNA-free TM kit (Invitrogen/ThermoFisher AM 1907) for 30 min at 37°C. RNA from liver (0.5µg) was reverse transcribed into cDNA using the Reverse Transcription System (Promega 3500). After 5-fold dilution, 5 µl was used as a template for real-time RT-PCR. Amplification was done for 40 cycles using SYBR^TM^ Green PCR master Mix (ThermoFisher, 4309155). Quantification of mRNA was performed using the ΔΔCT method and normalized to GAPDH for liver. Primer sequences are as follows: GAPDH (Accession number NM_008084) 5’-CTC ATG ACC ACA GTC CAT GCC A-3’; 5’-GGA TGA CCT TGC CCA CAG CCT T-3’; SAA1.1/2/1 (Accession number NM_009117.3) 5’-CTC CTA TTA GCT CAG TAG GTT GTG-3’; 5’-CAC TTC CAA GTT TCT GTT TAT TAC CC-3; SAA3 (Accession number NM_011315.3) 5’-TTT CTC TTC CTG TTG TTC CCA GTC-3’; 5’-TCA CAA GTA TTT ATT CAG CAC ATT GGG A-3’. For specific mRNAs, the primer sequences used in this study were as follows: SAA1.1 (accession no., NM009117), sense 5’–ATG AAG GAA GCT AAC TGG AAA AAC TC-3’, antisense 5’–TCC TCC TCA AGC AGT TAC TAC TGC AA-3’; SAA2.1 (accession no., NM011314), sense 5’–ATG AAG GAA GCT GGC TGG AAA GATGG-3’, antisense 5’–TCC TCC TCA AGC AGT TAC TAC TGC TC-3’ ; SAA3 (accession no., NM 011315), sense 5’–GCC ACC ATG AAG CCT TCC ATT GCC ATC ATT-3’.

### Cytokine Analysis

For quantitative analysis of cytokines, ELISA was used to measure the levels of tumor necrosis factor-α (TNF-α), interleukin-1β (IL-1β), or transforming growth factor-β (TGF-β) in brain tissue^67^. Cytokines were extracted from mouse brains as follows: frozen hemibrains were placed in tissue homogenization buffer containing protease inhibitor cocktail (Sigma, St Louis, MO, USA) 1:1000 dilution immediately before use, and homogenized using polytron. Tissue sample suspensions were distributed in aliquots and snap frozen in liquid nitrogen for later measurements. Invitrogen/ThermoFisher ELISA kits were then used, according to manufacturer directions (Carlsbad, CA, USA).

### Western blot assay

Coronal slices were dissected out after the indicated reperfusion time and the right hemisphere (the infarct/ischemic side) was selected. For western blotting, proteins were extracted from mouse brains using lysis buffer (50 mM Tris–HCl, pH 7.4, 0.5% Triton X-100, 4 mM EGTA, 10 mM EDTA, 1 mM Na_3_VO_4_, 40 mM Na_2_P_2_O_7_ꞏ10H_2_O, 50 mM NaF, 100 nM calyculin A, 50 μg/mL leupeptin, 25 μg/mL pepstatin A, 50 μg/mL trypsin inhibitor, and 1 mM dithiothreitol) as previously described^73^. The protein concentrations were quantified using Bradford’s assay and normalized. Equal quantities of protein were loaded and separated by 4-12% SDS–polyacrylamide gels (SDS/PAGE) and transferred to polyvinylidene difluoride (PVDF) or nitrocellulose membranes. Membranes were probed with primary antibodies overnight at 4°C and then with appropriate secondary antibodies. Thereafter, membranes were detected with an enhanced chemiluminescence (ECL) immunoblotting detection system (Amersham Biosciences, NJ, USA) using Luminescent Image Analyzer (LAS-4000 mini, Fuji Film, Tokyo, Japan). The densities of the bands were analyzed with ImageJ software (NIH, Bethesda, MA, USA). Primary antibodies were: SAA (R&D systems, AF2948); transferrin (abcam, ab84036); NLRP3 (abcam, ab214185); actin (abcam, ab179467). Secondary antibodies were: Goat anti-rabbit (Sigma, A6154).

### BBB Leakage Analysis

Evans blue (Sigma, CA, 2% in saline; 4 mL/kg) was used to determine the BBB permeability^74, 75^. For Evans blue detection, the dye was intravenously administered through tail vein 30 minutes prior to euthanization. Animals were transcardially perfused with saline, brains were sectioned with a 2 mm brain matrix, and images were taken by a photo scanner. Hemisphere samples were weighed, homogenized with 400 uL PBS, and precipitated with 50% trichloroacetic acid (Sigma, CA) overnight. All samples were centrifuged at 1,000 rpm for 30 minutes to separate out the brain tissue in pellet prior to measuring. Absorption was quantified at 610 nm with a plate reader (BMG Labtech, Cary, NC). To quantify Evans Blue penetration through the brain barrier, an Evans blue standardized curve was used. Results were quantified as microgram/ gram brain tissue.

### Adenoviral preparation

The production of replication defective adenoviral vectors expressing mouse SAA proteins has been described using second generation adenoviral vector^76, 77^. Briefly, Recombinant adenoviruses were expanded in 293 human embryonic kidney (HEK) cells. Virus was used to infect 293 cells and viral particles were isolated from the cells after 72 h of infection and purified by density gradient cesium chloride centrifugation. Virus particles were centrifuged through a two-step gradient of cesium chloride (1.45 g/ml and 1.20 g/ ml) for 2 h and 14 h, respectively. The virus particles were collected and chromatographed on a 10-ml Econo-Pac DG column as described by the manufacturer (Bio-Rad, Hercules, CA). The number of particles was determined by optical density.

### Statistics and Reproducibility

Statistical differences between parametric data (infarct volumes, activity values, ELISA values, cell counts, densitometry, and RNA data) were assessed using one-way analysis of variance (ANOVA) test with Bonferroni’s multi-group comparison, and non-parametric data (neurological deficits) were compared with the Kruskal-Wallis test with Dunn’s comparison (Prism 7.0, GraphPad). Sample sizes and number of replicates are included in the Figure legends or text. Survival was compared from all mice subjected to MCAo using the Kaplan-Meier test. Differences between data were considered statistically significant when *p* < 0.05.

